# Common germline polymorphisms and somatic cancer mutations exhibit non-random positional overlap across the human genome

**DOI:** 10.64898/2026.08.03.742387

**Authors:** Thais Silva Tavares, Daniela Souza Lima Barbosa, Renan Pedra de Souza, Roger Gomes da Silva, Thiago Peixoto Leal, Maycon Douglas de Oliveira, Carolina Silva de Carvalho, Luigi Marchionni, Mateus Henrique Gouveia, Francisco Pereira Lobo

**Affiliations:** Program of Bioinformatics, Institute of Biological Sciences, Federal University of Minas Gerais, Belo Horizonte, MG, Brazil; Department of Public Health Education, Morehouse School of Medicine, Atlanta, GA, USA; Pontifícia Universidade Católica de Minas Gerais PUC, Belo Horizonte, MG, Brazil; CIMO, LA SusTEC, Instituto Politécnico de Bragança, Campus de Santa Apolónia, Bragança, Portugal; Department of Genetics, Ecology and Evolution, Institute of Biological Sciences, Federal University of Minas Gerais, Belo Horizonte, MG, Brazil; Department of Medicine, Cancer Research Program, Centre for Translational Biology, McGill University Health Centre, McGill University, Montreal, QC, Canada; Department of Pathology and Laboratory Medicine, Weill Cornell Medicine, New York, NY, USA; Center for Cancer Research, National Cancer Institute, National Institutes of Health, Frederick, MD, USA

## Abstract

Germline and somatic mutations have traditionally been studied independently because they arise in distinct biological contexts and are shaped by different selective pressures. Despite these differences, both originate from the same molecular processes of DNA damage, replication error, and DNA repair. Yet this separation has limited the opportunity of investigation of genomic loci recurrently mutated across both mutational landscapes. Identifying such mutational co-occurrences may provide unique insights into the principles governing recurrent mutation. Here, we show that common germline polymorphisms and cancer-associated somatic SNVs recur at identical genomic positions across the human genome, sharing the same nucleotide substitutions more frequently than expected by chance. This recurrence persists within coding regions, is only minimally explained by the canonical hotspot contexts evaluated here (CpG islands and microsatellites), and is associated with a mutational signature profile enriched for the ubiquitous clock-like SBS5 signature together with DNA repair-associated signatures. Importantly, this overlap pattern is not shared across other germline variation: rare (AF<1%) and clinically classified variants exhibit significantly less overlap than expected. Together, these findings support the existence of intrinsically vulnerable genomic loci and provide a framework for investigating the mechanisms underlying recurrent mutation.

## INTRODUCTION

Mutations are the primary source of genetic variation and underlie both evolutionary change and human disease ^1,2^. Germline and somatic mutations arise in distinct biological contexts and are shaped by different selective pressures ^3–5^. Nevertheless, both ultimately originate from the fundamental molecular processes of DNA damage, replication error, and DNA repair ^6^. Understanding how these processes interact with locus-specific genomic properties is therefore central to explaining the heterogeneous distribution of mutations across the human genome.

Single nucleotide variants (SNVs) are the most abundant form of genetic variation in the human genome ^7^. Germline SNVs contribute to phenotypic diversity, adaptation, and hereditary diseases ranging from classical Mendelian disorders, such as sickle cell anemia ^8^ and cystic fibrosis ^9^, to complex polygenic traits ^10,11^. In contrast, somatic SNVs accumulate during an individual’s lifetime and play central roles in cancer initiation and progression ^12,13^. Among germline variants, common single nucleotide polymorphisms (SNPs), defined as variants with allele frequencies exceeding 1%, are generally considered evolutionarily tolerated or nearly neutral, although some contribute to adaptation ^14,15^. Conversely, deleterious variants are expected to remain rare because of purifying selection ^16^. The increasing availability of large-scale population, clinical, and cancer sequencing resources has greatly expanded our ability to investigate the diversity and distribution of these distinct classes of SNVs across the human genome ^17–21^.

Although individual mutation events occur stochastically, their distribution across the genome is not uniform. Certain genomic regions exhibit elevated mutation rates and are commonly referred to as mutational hotspots ^22^. Several factors influence this regional variation, including transcription-coupled repair, epigenetic modifications, DNA replication dynamics, and chromatin structure ^23^. Classic examples of mutational hotspots include CpG islands and microsatellites that are particularly susceptible to replication errors or repair defects ^24^. These regions are often associated with increased mutational susceptibility and have been implicated in multiple genetic diseases, including cancer ^25,26^.

Despite extensive research on mutational processes, relatively few studies have examined the spatial distribution of distinct SNV classes across the human genome from an integrative evolutionary and functional perspective ^24^. Most genomic studies focus on a single mutation type or disease context, such as cancer or hereditary disorders, resulting in fragmented views of mutational patterns. However, growing genomic resources now enable integrative approaches that compare mutation classes across biological contexts ^21^.

Recent studies have shown that germline variants and somatic cancer mutations can recur at the same genomic loci, despite addressing this overlap using distinct datasets and from different biological questions ^3,27–29^. Collectively, these observations suggest that recurrent positional overlap may reflect mutational vulnerabilities shared across germline and somatic contexts. However, no previous study has systematically determined whether this phenomenon is broadly distributed across germline variation or preferentially concentrated within specific variant classes. Here, we address this question through the first unified genome-wide comparison of recurrent germline-somatic mutation across common, rare, benign, and pathogenic germline variants. Using population variants from the Human Genome Diversity Project (HGDP) ^20^ and clinically annotated variants from ClinVar ^18,19^, together with somatic cancer mutations from the Catalogue Of Somatic Mutations In Cancer (COSMIC) ^30,31^, we tested whether exact-coordinate overlaps occurred more frequently than expected by chance. Common germline polymorphisms were the only variant class significantly enriched for recurrent positional overlap, both genome-wide and within coding sequences. This class-specific pattern was independently replicated using the high-coverage 1000 Genomes Project (1KGP) call set ^32,33^, whereas rare variants were depleted in both population panels and ClinVar benign and pathogenic variants were depleted within the harmonized coding-sequence analysis.

We next investigated the sequence, functional, and mutational characteristics of these recurrent loci, including their distribution across coding regions, canonical mutational hotspots (CpG islands and microsatellites), mutational signatures, and biological pathways. Recurrent positional overlap remained enriched within coding regions and was only minimally explained by canonical mutational hotspots. Shared coding loci were enriched for clock-like single-base substitution (SBS) ^34–36^ mutational signatures, especially SBS5, a clock-like mutational signature of unknown etiology, together with signatures associated with defective DNA repair and genome instability. Functional enrichment further linked these loci to cancer-associated pathways, including PI3K-AKT signaling and extracellular matrix-receptor interaction. Shared coding loci were characterized for SBS mutational signatures by a prominent contribution of SBS5 together with repair and genome instability-associated signatures (SBS1, SBS6, SBS54, and SBS87). Together, these findings reveal a previously unrecognized positional relationship between germline and somatic mutational landscapes and establish a foundation for investigating why certain genomic loci recurrently accumulate mutations across biological contexts.

## RESULTS

### Dataset composition and genomic distribution of germline and somatic SNV sets

We first characterized the composition and genomic distribution of the germline and somatic SNV datasets included in this study. The primary analyses integrated three genomic resources and generated five SNV sets representing distinct biological classes: COSMIC somatic SNVs, HGDP common and rare germline SNVs, and ClinVar benign/likely benign and pathogenic/likely pathogenic germline SNVs (**Supplementary Table S1**). For independent replication of the population-based genome-wide findings, high-coverage 1KGP variants were processed using the same analytical pipeline and classified as common or rare according to the same allele-frequency criteria.

Genomic annotation using the Ensembl Variant Effect Predictor (VEP) ^37^ revealed marked differences in the distribution of SNVs across genomic compartments. HGDP common and rare SNVs and COSMIC mutations were predominantly located outside coding sequences, whereas ClinVar benign and pathogenic SNVs were concentrated within CDS, consistent with the coding-focused ascertainment of clinically interpreted variation (**Fig. 1A**). Because this imbalance in coding-region representation could confound genome-wide overlap estimates, ClinVar datasets were not included in whole-genome overlap tests and were evaluated exclusively in CDS-restricted comparisons. For these analyses, all five primary SNV sets were restricted to MANE-defined coding sequences (Matched Annotation from NCBI and EMBL-EBI) ^38^, thereby establishing a common genomic search space across variant classes (**Fig. 1B**).

**Figure 1.**
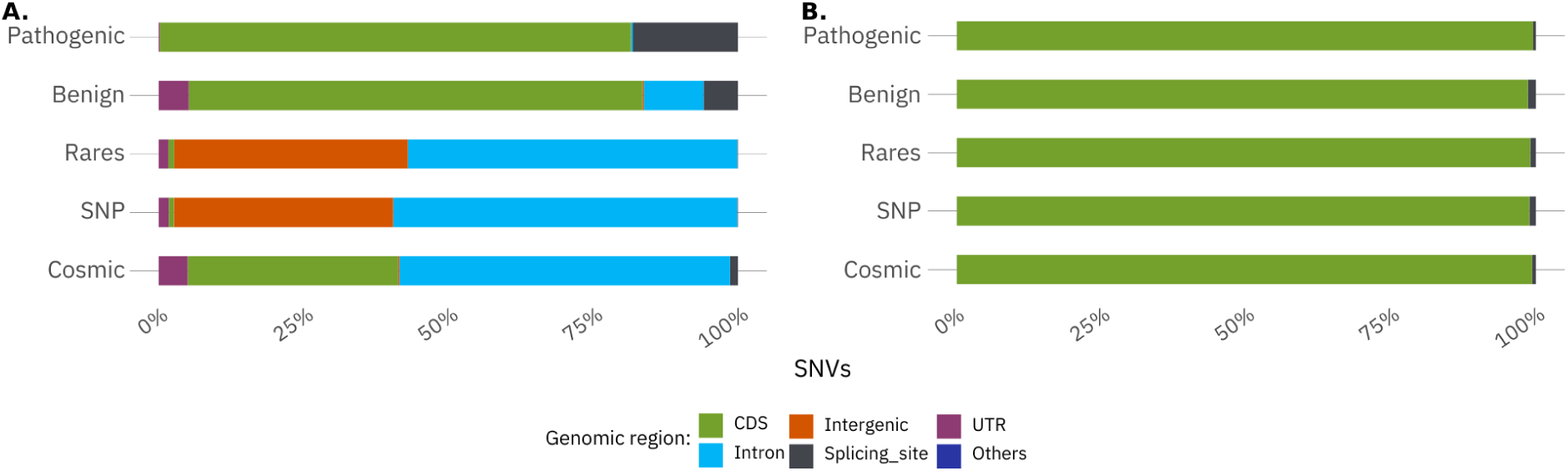
Genomic distribution of SNVs across the five SNV datasets. (**A**) WG distribution of SNVs across genomic regions. Population-based datasets (HGDP common and rare SNVs) and COSMIC mutations are predominantly located in non-coding regions, whereas ClinVar benign and pathogenic SNVs are strongly enriched in coding sequences. (**B**) Distribution after restricting all datasets to CDS, illustrating the harmonized genomic composition used in downstream CDS-restricted analyses. Pearson’s chi-square test indicated significant differences in genomic region composition among WG datasets (χ^2^ = 21,977.078; P < 2.2 × 10^-16^).

Functional consequence annotations within CDS preserved the expected distinctions among variant classes. ClinVar pathogenic SNVs showed the highest proportions of stop-gained and high-impact annotations, whereas ClinVar benign SNVs were predominantly synonymous and classified as low impact. HGDP common and rare SNVs and COSMIC mutations were dominated by missense and synonymous consequences, although their relative proportions differed across datasets (**Supplementary Fig. S1**). Thus, restriction to MANE-defined CDS harmonized the genomic background used for overlap testing while preserving the distinct functional profiles of the analyzed SNV classes.

### Genome-wide overlap reveals enrichment exclusively for common germline SNVs

We next tested whether COSMIC somatic SNVs overlapped germline SNVs at identical genomic whole-genome (WG) coordinates more frequently than expected by chance in each population reference panel using Fisher’s exact tests.

WG analysis using the HGDP reference panel revealed significant enrichment of exact-coordinate overlaps between common germline SNPs and COSMIC mutations (OR = 1.920, 95% CI = 1.904-1.936, P < 2.2 × 10^-16^; **Fig. 2 and Supplementary Table S2**). In contrast, rare germline SNVs were significantly depleted relative to expectation (OR = 0.668, 95% CI = 0.664-0.671, P < 2.2 × 10^-16^), demonstrating that recurrent positional overlap with somatic mutations is not a general property of germline variation.

**Figure 2.**
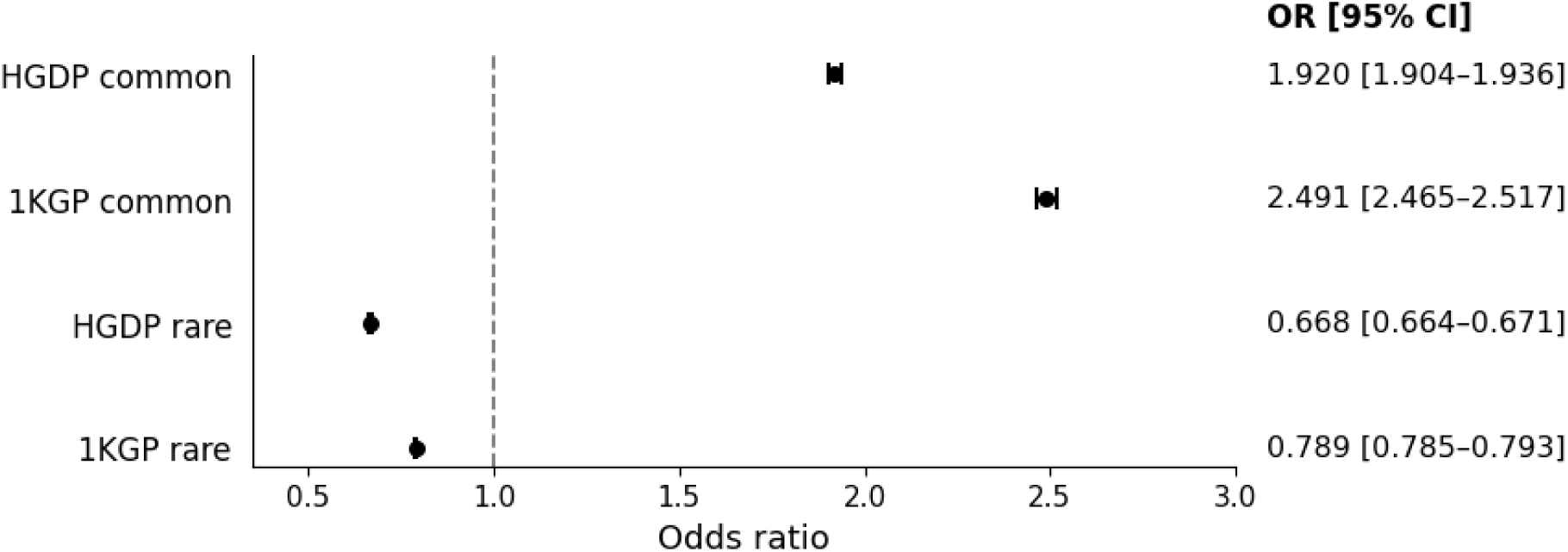
Genome-wide enrichment of exact-coordinate overlap between COSMIC somatic mutations and population-based germline SNV datasets. Forest plot showing odds ratios (ORs) and 95% confidence intervals (CIs) from Fisher’s exact tests assessing exact-coordinate overlap between COSMIC somatic SNVs and LD-pruned common and rare germline SNVs from the HGDP and high-coverage 1000 Genomes Project (1KGP) reference panels. The dashed vertical line indicates the null value (OR = 1), and points and horizontal bars represent ORs and 95% CIs, respectively. Common germline polymorphisms were significantly enriched for overlap with COSMIC mutations in both reference panels, whereas rare germline SNVs were consistently depleted. Complete statistical results are provided in Supplementary Table S2.

To assess the robustness of this pattern to the choice of population reference panel, we repeated the genome-wide analysis using linkage disequilibrium (LD)-pruned SNVs from the high-coverage 1KGP dataset. The direction of the associations was consistently reproduced: exact-coordinate overlap between 1KGP common polymorphisms and COSMIC mutations was significantly enriched (OR = 2.491, 95% CI = 2.465–2.517, P < 2.2 × 10^-16^; **Fig. 2** and **Supplementary Table S2**), whereas overlap involving rare 1KGP SNVs was significantly depleted (OR = 0.789, 95% CI = 0.785–0.793, P < 2.2 × 10^-16^). The concordant results across HGDP and 1KGP demonstrate that the class-dependent pattern is robust to the choice of population reference panel and argue against a panel-specific artifact.

Together, these analyses establish a reproducible frequency-dependent pattern of germline-somatic recurrence: exact-coordinate overlap with COSMIC mutations is enriched among common germline polymorphisms but depleted among rare germline variants. This contrast motivated subsequent analyses of the functional characteristics, mutational spectra, and biological pathways associated with shared germline-somatic loci.

To complement the Fisher’s test results and determine whether the primary genome-wide enrichment signal could arise under random genomic positioning, we performed a chromosome-constrained permutation analysis of the HGDP common SNP dataset. HGDP SNP positions were randomized while preserving chromosome assignment and the number of SNVs per chromosome. Across 1,000 permutations, the mean expected overlap with COSMIC mutations was 7,943.5 loci (empirical 95% interval: 7,773-8,130). The observed overlap of 59,569 loci was 7.50-fold greater than the null mean, and none of the permuted datasets produced an overlap equal to or greater than the observed value. Using a one-sided plus-one corrected estimate, this corresponded to an empirical P = 9.99 × 10^-4^ (**Fig. 3**). This analysis was applied to the HGDP comparison because it constituted the primary discovery analysis, whereas 1KGP was used to evaluate replication across an independent population reference panel.

**Figure 3.**
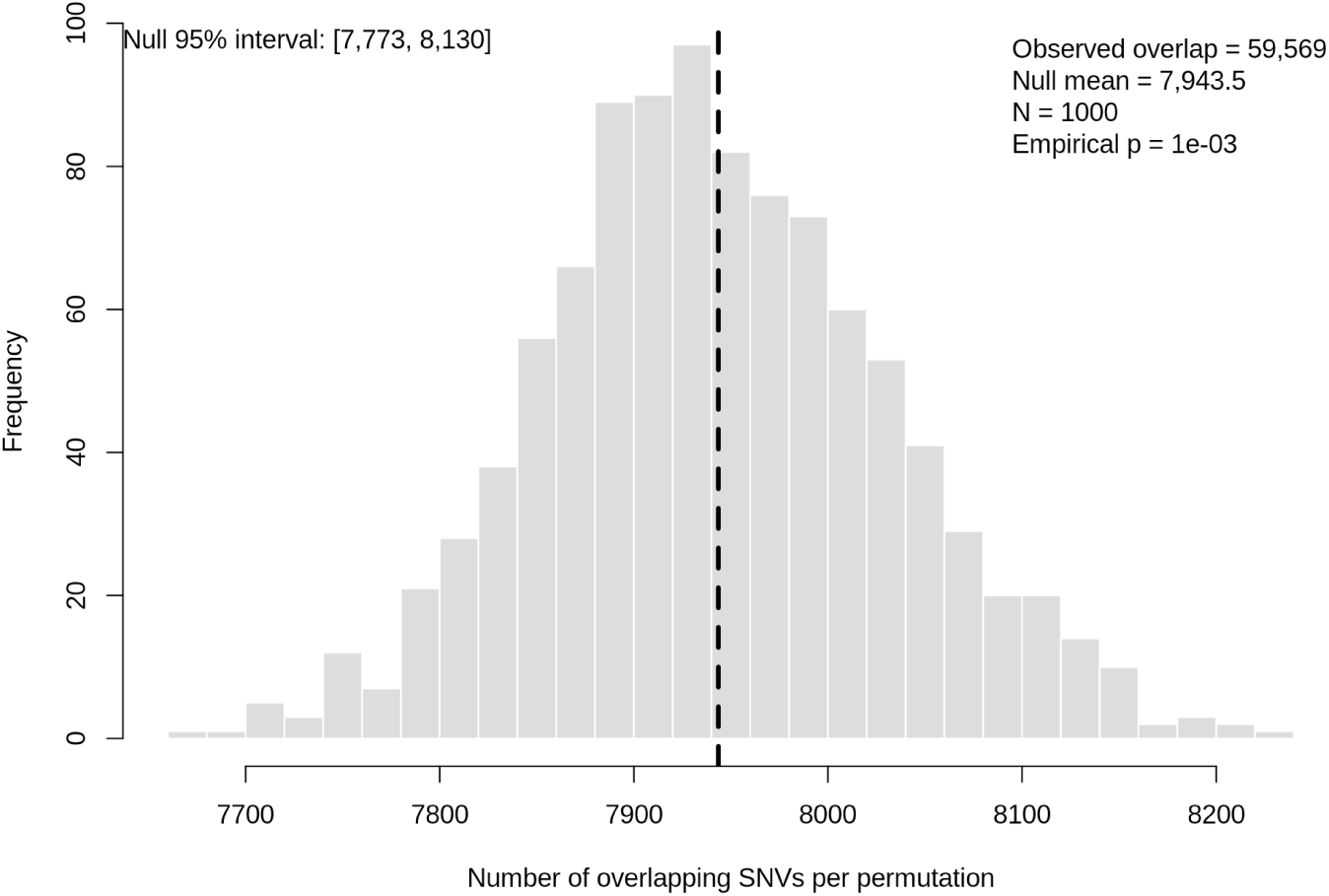
Permutation-based validation of the genome-wide overlap between HGDP common polymorphisms and COSMIC somatic mutations. HGDP common SNP positions were randomly shuffled within chromosomes while preserving chromosome assignment and per-chromosome SNV counts, and exact-coordinate overlaps with COSMIC mutations were recalculated across 1,000 permutations. Grey bars show the empirical null distribution, and the dashed vertical line indicates its mean (7,943.5 overlaps; empirical 95% interval: 7,773-8,130). The observed overlap of 59,569 loci was 7.50-fold greater than the null mean and exceeded the overlap obtained in every permutation. Because the observed value lies outside the displayed range of the null distribution, it is indicated as off-scale. The one-sided empirical P value was calculated using the plus-one correction and was 9.99 × 10^-4^.

We next examined whether genome-wide positional overlaps also involved identical nucleotide substitutions. At each shared coordinate, all observed REF>ALT substitutions were retained, and a locus was considered concordant when at least one identical substitution was present in both the germline and somatic datasets. Among the 59,569 loci shared between HGDP common SNPs and COSMIC mutations, 56,932 (95.57%) contained an identical REF>ALT substitution, whereas 2,637 (4.43%) showed positional overlap without an exact substitution match (**Table 1**). Exact-substitution concordance was also observed at the majority of loci shared between rare SNVs and COSMIC mutations (106,477 of 135,458; 78.61%), although at a lower proportion than among common SNPs; 28,981 loci (21.39%) lacked an identical substitution. Thus, despite their contrasting positional enrichment patterns, overlaps in both germline frequency classes predominantly represented recurrence of the same nucleotide change, with markedly higher allele-level concordance at common SNP loci.

**Table 1.** Genome-wide nucleotide-substitution concordance at exact-coordinate overlaps between HGDP germline SNVs and COSMIC somatic mutations. A locus was considered concordant when at least one identical REF>ALT substitution was present in both datasets. Each genomic coordinate was counted once, and all substitutions were retained at multiallelic loci.

| Comparison | Overlap | ALT same | ALT different |
| --- | --- | --- | --- |
| Common SNPs × COSMIC | 59,569 | 56,932 (95.57%) | 2,637 (4.43%) |
| Rare SNVs × COSMIC | 135,458 | 106,477 (78.61%) | 28,981 (21.39%) |

### Coding-restricted analyses reveal enrichment specific to common germline SNVs

Restricting analyses to MANE-defined coding sequences (CDS) confirmed that germline-somatic positional overlap observed in the WG analysis was retained within coding sequences. Within CDS, only common germline SNVs showed significant enrichment with COSMIC mutations (OR = 5.65), whereas all other COSMIC comparisons were depleted, including rare SNVs (OR = 0.52), ClinVar benign SNVs (OR = 0.70), and ClinVar pathogenic SNVs (OR = 0.56; **Figure 4**; **Supplementary Table S2**).

**Figure 4.**
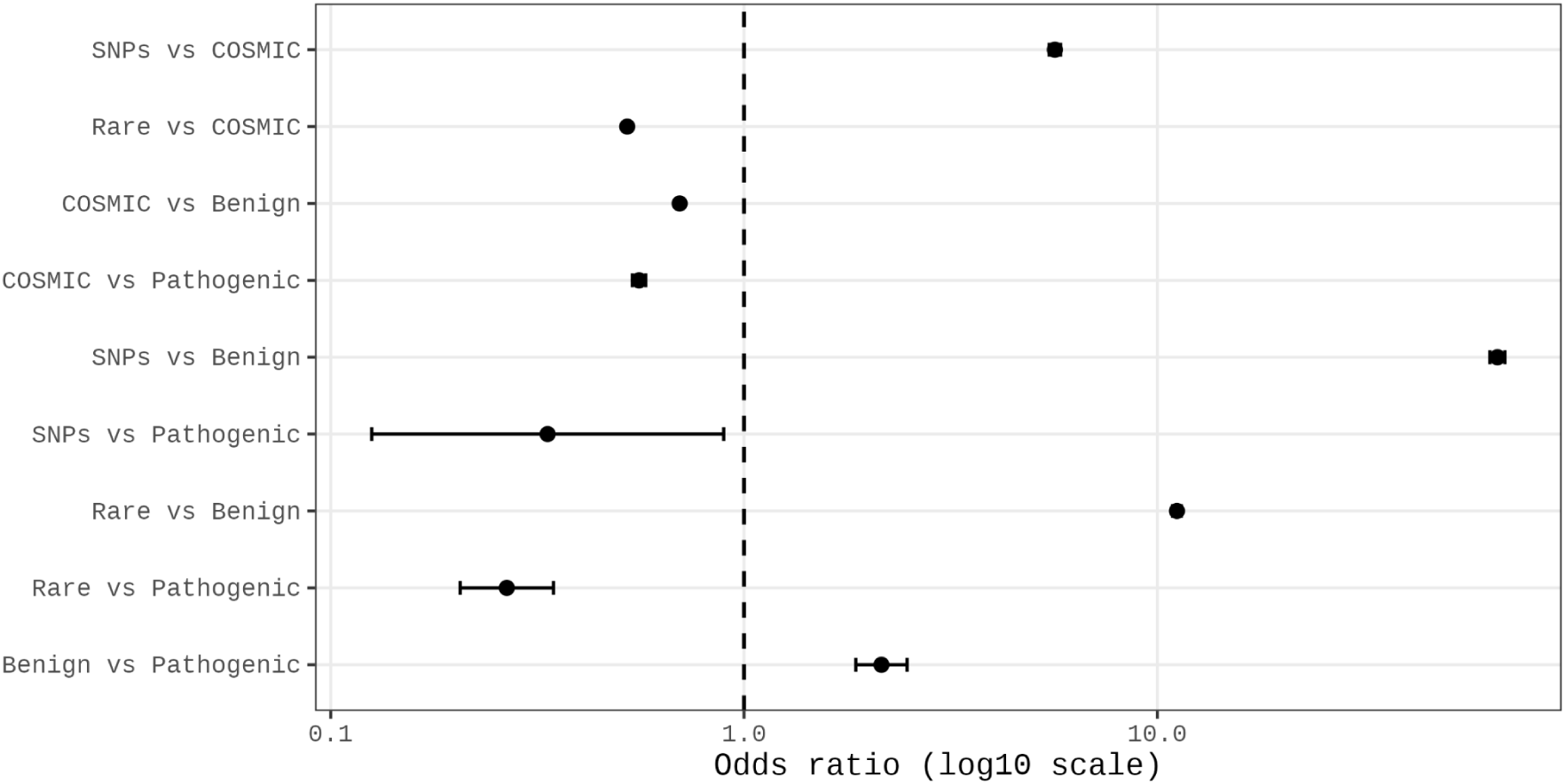
Coding-sequence overlap enrichment across germline and somatic SNV classes. Odds ratios (OR, log scale) from Fisher’s exact tests assessing exact-coordinate overlap between SNV classes restricted to MANE-defined coding sequences (CDS). The dashed vertical line indicates no enrichment (OR = 1). Common germline SNPs exhibit strong enrichment of shared loci with COSMIC somatic mutations, whereas rare germline SNVs and clinically annotated SNVs (ClinVar benign and pathogenic) show significant depletion relative to COSMIC. Error bars represent 95% confidence intervals. Complete statistical results are provided in **Supplementary Table S2.**

Pairwise comparisons among germline datasets revealed additional class-specific patterns. Both common and rare SNVs were enriched for positional overlap with ClinVar benign variants, whereas ClinVar pathogenic variants were depleted relative to both population-based germline classes. Benign and pathogenic ClinVar variants nevertheless exhibited positive positional overlap. These contrasting results indicate that positional overlap within CDS depends on the specific variant classes being compared and cannot be explained solely by their shared localization within coding sequences. Although these patterns may be compatible with differences in functional constraint among variant classes, evolutionary constraint was not directly evaluated in the present analysis.

Together, the CDS-restricted analyses showed that the enrichment of common germline-somatic overlap identified in the whole-genome analysis was retained within MANE-defined coding sequences. Common SNVs were the only germline class enriched for positional overlap with COSMIC mutations, whereas rare SNVs and ClinVar benign and pathogenic SNVs were depleted. Thus, the enrichment observed at common germline loci cannot be attributed solely to the overall genomic distribution of the variant datasets or to their localization within coding sequences.

### Mutational signature profiles reveal a prominent SBS5 contribution at shared germline-somatic coding loci

To characterize the nucleotide-substitution patterns associated with germline-somatic shared loci, we generated a pooled SBS96 catalogue of COSMIC substitutions occurring at coding positions shared with common germline SNVs. The observed catalogue was decomposed into COSMIC v3.4 reference signatures using SigProfilerAssignment ^34–36^, as described in the Methods. CDS decomposition identified SBS5 as the largest contributor to the reconstructed mutational spectrum (37.3%), followed by SBS87 (21.5%), SBS54 (15.3%), SBS6 (13.0%), and SBS1 (12.9%) (**Figure 5**). The reconstructed spectrum closely reproduced the observed trinucleotide profile (cosine similarity = 0.980; correlation = 0.971), indicating that the assigned combination of reference signatures captured the major features of the mutation catalogue. Thus, shared coding loci exhibited a heterogeneous mutational spectrum characterized by a prominent SBS5 contribution together with four additional SBS signatures.

**Figure 5.**
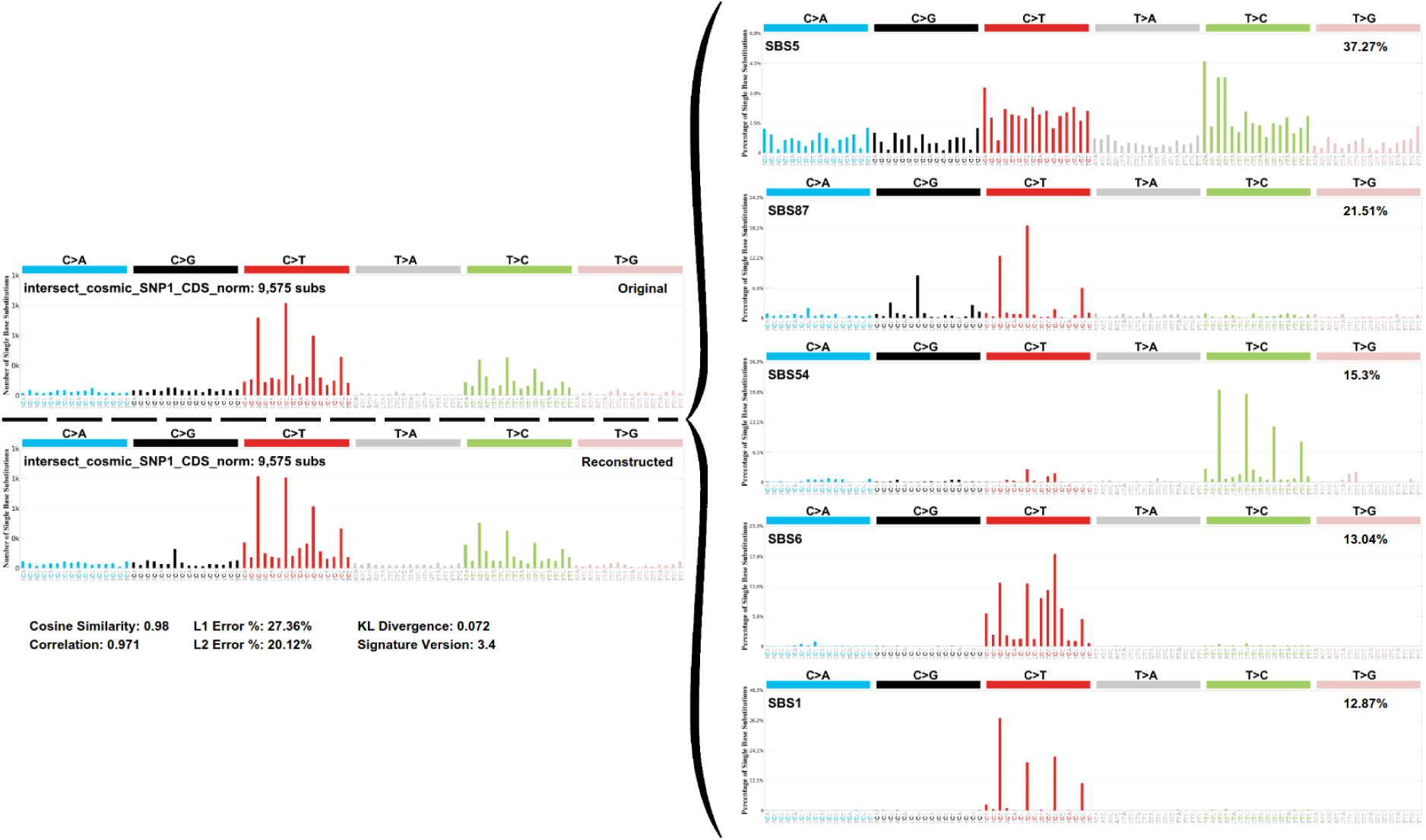
Mutational signature decomposition of COSMIC substitutions at coding loci shared with common germline SNVs. Single-base substitution (SBS96) signature analysis was performed on COSMIC mutations occurring at loci shared with common germline SNVs within MANE-defined coding sequences. The left panels show the observed and reconstructed SBS96 spectra, which exhibited a cosine similarity of 0.98. The right panels show the contributions of SBS5 (37.27%), SBS87 (21.51%), SBS54 (15.30%), SBS6 (13.04%), and SBS1 (12.87%) to the reconstructed spectrum.

Because SBS1 is associated with the deamination of 5-methylcytosine at CpG dinucleotides and SBS6 is associated with defective DNA mismatch repair and microsatellite instability ^35^, we next evaluated whether canonical context-dependent mutational features contributed to the observed germline-somatic overlap. Both CpG islands and microsatellite regions were significantly enriched among shared loci, with odds ratios of 7.49 and 3.85, respectively (**Supplementary Table S3**). Nevertheless, CpG islands intersected 2,839 of the 59,569 shared loci (4.77%), whereas microsatellites intersected only 131 loci (0.22%). Thus, although these genomic contexts are preferentially represented among shared loci, they encompass only small proportions of the overall positional overlap.

### Genes harboring shared coding loci converge on interconnected signaling and extracellular matrix pathways

To determine whether exact-coordinate recurrence also converged at the gene and pathway levels, we analyzed 6,149 unique genes harboring coding loci shared between common germline SNPs and COSMIC mutations. Kyoto Encyclopedia of Genes and Genomes (KEGG) over-representation analysis identified 10 significantly enriched pathways (FDR < 0.05), comprising 511 unique mapped genes and 740 gene-pathway associations (**Figure 6, Supplementary Table S4**). ECM-receptor interaction exhibited the highest enrichment ratio, whereas PI3K-AKT signaling contained the largest number of mapped genes (n = 143). Focal adhesion and ECM-receptor interaction contained 99 and 61 mapped genes, respectively.

**Figure 6.**
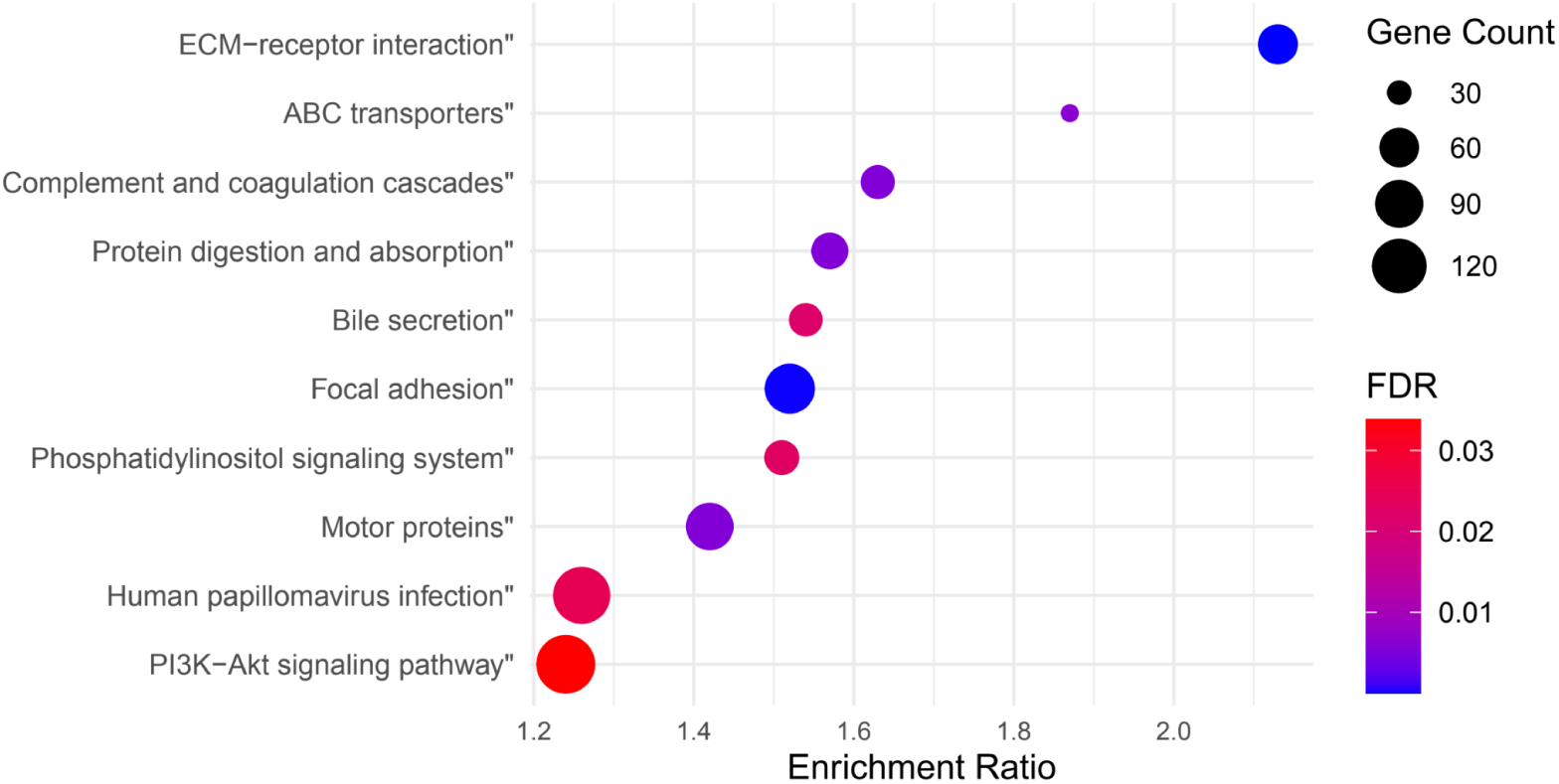
Functional enrichment of genes intersecting shared coding germline-somatic loci. KEGG over-representation analysis of genes harboring coding loci shared between common germline SNPs and COSMIC somatic mutations. Enriched pathways include cancer-associated signaling networks and extracellular matrix-related processes. Dot size represents the number of genes per pathway, and color indicates statistical significance after multiple-testing correction.

Substantial gene overlap was observed among PI3K-AKT signaling, ECM-receptor interaction, and focal adhesion, with 48 genes annotated to all three pathways. This shared subset included 13 collagen genes, 12 integrin genes, and 10 laminin genes, together with FN1, TNC, THBS1, and THBS2. PIK3R1 and PIK3R2 further connected PI3K-AKT and focal adhesion with the phosphatidylinositol signaling system. Thus, the significant pathway enrichments reflected convergence across multiple interconnected genes and gene families rather than isolated gene-level signals.

A complementary over-representation analysis based on cytogenetic-band annotations identified significant enrichment of genes harboring shared loci across six genomic regions: 13q12.12, 19p13.12, 19p13.3, 22q13.33, 5q31.3, and 6q27 (FDR < 0.05; Supplementary **Figure S2, Supplementary Table S5**). These regions comprised 230 unique genes, with the largest mapped gene set observed at 19p13.3 (n = 92), followed by 5q31.3 (n = 53) and 19p13.12 (n = 33). The analysis additionally returned the GRCh38 assembly patch HG2308_PATCH, containing 23 genes that were also assigned to 5q31.3; this patch was therefore considered an alternative assembly annotation rather than an independent enriched genomic region.

Together, these analyses indicate that genes harboring shared germline-somatic coding loci are not uniformly distributed across functional or cytogenetic annotations but converge on an interconnected ECM-integrin-PI3K signaling axis and on discrete genomic regions, including two adjacent bands on chromosome 19p. This combined functional and regional pattern supports a non-random organization of germline-somatic recurrence and prioritizes these pathways and genomic regions for investigating the mechanisms underlying positional overlap.

## DISCUSSION

In this study, we integrated population-based (HGDP)^20^, clinically annotated (ClinVar)^18,19^, and cancer-associated (COSMIC)^30,31^ SNV datasets to determine whether recurrent positional overlap with somatic mutations differs across germline variant categories. Our central finding is that germline-somatic recurrence is not a general property of germline variation but is preferentially concentrated at common polymorphic sites. Common SNVs showed significant enrichment for exact-coordinate overlap with COSMIC mutations in the HGDP, a pattern independently replicated using the 1KGP^32,33^ and retained within coding sequences. Moreover, the observed genome-wide overlap exceeded the chromosome-preserving permutation expectation by more than sevenfold, demonstrating that the signal cannot be explained by random genomic positioning alone.

The contrasting behavior of common and rare germline SNVs provides an evolutionary context for this pattern. Common polymorphisms have persisted across population history and are generally more compatible with evolutionary tolerance, whereas rare variants include a larger proportion of recently arisen and potentially deleterious alleles subject to purifying selection ^2,39^. Nevertheless, population frequency cannot be interpreted as a direct measure of neutrality, because allele frequencies are also shaped by demographic history, genetic drift, local mutation rates, and positive selection. Our findings therefore reveal a frequency-dependent pattern of germline-somatic recurrence: positional overlap with somatic cancer mutations was enriched among common germline polymorphisms, whereas rare germline variants were consistently depleted. This contrast may reflect contributions from evolutionary constraint, locus-specific mutational processes, or both, although our analyses did not directly disentangle these mechanisms. This interpretation is consistent with evidence that mutation rates vary across the human genome and that variants arising at recurrently mutated sites are subsequently shaped by selection ^2,24^.

Our findings place previous observations of germline-somatic overlap into a unified comparative framework. Lu et al. identified rare germline variants at recurrent somatic mutation sites in the context of cancer predisposition ^27^, whereas Meyerson et al. showed that identical variants shared between non-common germline and somatic cancer datasets occur more frequently than expected and are largely attributable to shared nucleotide-context-dependent mutational vulnerabilities ^3^. Conversely, Wang et al. demonstrated a 2.7-fold higher incidence of somatic mutations at common polymorphic loci, consistent with the enrichment observed here ^28^. Exact matches between population variants and COSMIC entries have also been interpreted as potential germline misclassification ^29^. Although some contribution from this source cannot be excluded, the restriction of enrichment to common polymorphisms, its replication across independent population panels, and its persistence within coding regions make nonspecific database overlap or germline contamination unlikely to explain the complete pattern. Consistently, within the harmonized coding-sequence background, rare, ClinVar benign, and ClinVar pathogenic variants were depleted relative to COSMIC mutations. Together, these findings support a reproducible, class-dependent pattern of germline-somatic recurrence that extends beyond technical overlap or clinical ascertainment.

Beyond positional enrichment, we examined whether germline and somatic SNVs overlapping at the same coordinates also shared the same alternate allele. Importantly, positional depletion did not imply an absence of shared loci: although rare germline SNVs overlapped COSMIC mutations less frequently than expected from the genomic background, a substantial number of rare germline-somatic overlaps remained. Among overlapping loci, alternate-allele concordance was high for both common and rare germline SNVs, indicating that overlap frequently involved the same nucleotide substitution rather than different substitutions at the same coordinate. However, concordance was substantially higher for common SNPs than for rare SNVs (95.57% versus 78.61%, respectively), indicating that the two germline categories should not be interpreted as exhibiting equivalent allele-matching patterns.

The coexistence of positional depletion and high allele concordance among rare SNVs highlights the distinction between the frequency and identity of overlap. Rare germline SNVs shared fewer loci with COSMIC than expected, but when overlap occurred, it frequently involved the same alternate allele. Common polymorphisms differed in both dimensions: their loci were enriched for overlap with COSMIC and showed particularly high allele concordance. Thus, positional enrichment measures how often loci are shared relative to the genomic background, whereas allele concordance determines whether the shared coordinate contains the same nucleotide substitution.

Mutational signature analysis characterized the substitution spectra at germline-somatic shared loci and highlighted candidate mutational processes associated with these positions. The CDS profile retained SBS5 as the largest contributor while revealing a broader combination of SBS1, SBS6, SBS54, and SBS87.

SBS5 is a ubiquitous clock-like signature whose burden generally increases with age across normal and cancer tissues, although its rate of accumulation varies among tissues and its underlying etiology remains incompletely understood ^35^. Its consistent prominence at shared loci raises the possibility that the still poorly understood processes captured by SBS5 contribute to the mutational spectrum at recurrent genomic positions, making this signature a compelling candidate for future mechanistic investigation. SBS1, which primarily reflects deamination of 5-methylcytosine at CpG dinucleotides, provided additional evidence of a broadly acting endogenous process. Together, SBS1 and SBS5 indicate that shared loci carry nucleotide substitution patterns compatible with ubiquitous, time-associated mutational processes. This interpretation complements the findings of Meyerson et al. ^3^, who showed that substitution class and extended nucleotide context explain a substantial proportion of identical variants shared between germline and somatic datasets.

The additional coding signatures point to more context-specific mutational influences. SBS6 is associated with defective DNA mismatch repair and microsatellite instability, whereas SBS87 has been experimentally linked to thiopurine treatment and may reflect treatment-related processes represented within the COSMIC records ^35,40^. SBS54 has a less certain biological interpretation, with possible contributions from sequencing artefacts or germline contamination, and should therefore be considered separately from repair-associated signatures ^30,35^. Independent genomic analyses further showed that CpG islands and microsatellites were significantly enriched among shared loci but accounted for only approximately 5% and 0.2% of shared events, respectively. Thus, canonical hotspot contexts contribute to positional overlap but explain only a small fraction of its genome-wide extent. Collectively, these findings support the contribution of multiple mutational influences, while the consistent predominance and unresolved biology of SBS5 provide a particularly important clue for understanding recurrence across germline and somatic contexts.

Moving beyond SNV-level recurrence, gene-based over-representation analysis showed that genes harboring coding loci shared between common germline SNPs and COSMIC mutations were enriched in cancer-relevant pathways, particularly PI3K-AKT signaling, ECM-receptor interaction, and focal adhesion. These pathways formed an interconnected functional axis supported by 48 genes annotated to all three pathways, prominently including collagen (COL), integrin (ITGA/ITGB), and laminin (LAMA/LAMB/LAMC) family members, together with FN1, TNC, THBS1, and THBS2. PIK3R1 and PIK3R2 further connected PI3K-AKT and focal adhesion with phosphatidylinositol signaling. The enrichment of PI3K-AKT signaling is biologically relevant because dysregulation of this pathway is widespread across human cancers and coordinates cellular survival, proliferation, and metabolic signaling ^41,42^. In parallel, ECM-receptor interactions and focal adhesions translate cell-matrix contacts into integrin and FAK-mediated signaling, connecting extracellular and mechanical cues to cytoskeletal organization, adhesion, and migration ^43,44^. Their joint enrichment therefore places shared germline-somatic loci at the interface between growth-factor signaling and microenvironmental regulation and prioritizes ECM-integrin-PI3K crosstalk for future functional investigation.

Cytogenetic-band enrichment provided complementary evidence that genes harboring shared coding loci are regionally concentrated. Among the six significantly enriched bands, two were located on the short arm of chromosome 19: 19p13.12 and 19p13.3, comprising 33 and 92 mapped genes, respectively. The 19p13.3 signal was particularly relevant because STK11, a negative regulator of mTOR signaling, contributed to both this cytogenetic enrichment and the enriched PI3K-AKT pathway, directly connecting the regional and functional results ^41,45^. Using a different analytical framework, Carter et al. previously found that germline variation at 19p13.3 was associated with a fourfold increased likelihood of somatic PTEN mutations and provided functional evidence linking this region to PTEN-dependent mTOR signaling ^46^. This independent observation reinforces 19p13.3 as a particularly relevant region for germline-somatic interactions. Together, the convergence of chromosome 19p enrichment with PI3K-AKT-mTOR signaling suggests that regional genomic organization and pathway context may jointly contribute to the recurrence of mutations across germline and somatic settings.

Several limitations should be acknowledged. First, our analyses were restricted to single-nucleotide substitutions and did not capture indels or structural variants. Second, although LD pruning and chromosome-preserving permutations reduced major sources of confounding, the null model did not explicitly match variants according to sequence context, mappability, replication timing, chromatin state, or other determinants of regional mutation opportunity. Residual effects of regional mutation-rate heterogeneity therefore cannot be completely excluded. Third, COSMIC integrates somatic variants across heterogeneous tumor types, study designs, and sequencing strategies, with extensive representation from targeted and whole-exome sequencing studies and a more recent expansion of whole-genome datasets ^30^. Consequently, genomic regions are not uniformly interrogated, and differences in coverage, tumor-type composition, matched-normal practices, and potential germline misclassification may influence overlap estimates, particularly in genome-wide analyses. The population and clinical databases used here also differ in ancestry composition, callability, and clinical ascertainment. Nevertheless, replication across HGDP and 1KGP, together with the contrasting behavior of common, rare, benign, and pathogenic variants, makes nonspecific database ascertainment unlikely to explain the complete class-dependent pattern. Fourth, mutational signatures were inferred from pooled COSMIC mutations at shared loci without comparison with a matched COSMIC background; they therefore characterize the mutational spectrum and nominate candidate processes but do not demonstrate signature-specific enrichment or establish that the corresponding germline variants arose through the same processes. Finally, functional enrichment depends on the annotation systems and reference backgrounds employed and identifies pathway-level convergence rather than pathway activation or functional effects of individual variants. Accordingly, the observational design cannot directly distinguish the relative contributions of local mutability, genomic architecture, selection, and technical ascertainment to positional recurrence.

In summary, we demonstrate that exact-coordinate overlap between germline and somatic cancer SNVs is a reproducible, class-dependent feature of the human genome, enriched specifically at common germline polymorphisms. This pattern replicated across independent population datasets, persisted within coding regions, and was only minimally explained by the canonical hotspot contexts examined here. Shared coding loci were further characterized by an SBS5-dominated mutational spectrum and functional convergence on interconnected ECM-integrin-PI3K networks and discrete cytogenetic regions. By providing the first systematic genome-wide comparison across biologically distinct classes of germline variation, our study establishes recurrent germline-somatic loci as a tractable set for investigating the sequence, genomic, and mutational features that make particular positions repeatedly susceptible to mutation across biological contexts.

## Supporting information

Supplementary Table S5. Complete list of genes present in the significantly enriched cytogenetic bands identified in the Over-Representation Analysis

Supplementary Table S4. Complete list of genes enriched in the KEGG pathways identified in the Over-Representation Analysis (ORA).

## Author contributions

T.S.T. conducted the investigation, curated the data, developed the software and methodology, performed the formal analyses and visualisation, interpreted the results, and wrote the original draft of the manuscript.

D.S.L.B., M.D.O., L.M., and R.G.S. contributed to data analysis, visualisation, interpretation of the results, and review and editing of the manuscript.

C.S.C. and T.P.L. contributed to the analysis of population-genetics data, interpretation of the results, and review and editing of the manuscript.

R.P.S. contributed to supervision, methodology development, data interpretation, visualisation, and review and editing of the manuscript.

M.H.G. contributed to supervision, conceptualisation, interpretation of the results from a population-genetics perspective, secured funding for publication, and reviewed and edited the manuscript.

F.P.L. conceived the study, supervised the project, contributed to methodology development and interpretation of the results, and reviewed and edited the manuscript.

All authors reviewed and approved the final version of the manuscript.

## Acknowledgements

The authors acknowledge the Graduate Program in Bioinformatics at the Universidade Federal de Minas Gerais (UFMG) for institutional support and the Centro de Processamento de Alto Desempenho of the Institute of Biological Sciences (CEPAD-ICB-UFMG) for providing computational resources through the Sagarana high-performance computing cluster.

## METHODS

### Study design and analytical overview

We performed a genome-wide integrative analysis to evaluate whether germline and somatic SNVs co-occur at identical genomic coordinates more frequently than expected by chance. The workflow included: (i) quality control and harmonization of SNV sets, (ii) linkage disequilibrium (LD) pruning of germline SNVs, (iii) exact-coordinate overlap testing using Fisher’s exact tests, (iv) permutation-based null modeling, and (v) functional characterization of shared loci through mutational signature and pathway enrichment analyses.

### SNV datasets

#### Germline SNVs

Population-level whole-genome germline SNVs were obtained from the HGDP ^20^, which served as the primary reference dataset. Independent validation of the genome-wide overlap analyses was subsequently performed using the high-coverage 1000 Genomes Project (1KGP) call set ^32,33^. Clinically annotated germline SNVs restricted to protein-coding sequences (MANE-defined ^38^ CDS) were obtained from ClinVar ^18,19^. ClinVar SNVs were categorized as benign/likely benign or pathogenic/likely pathogenic according to clinical assertions. Pathogenic SNVs were further restricted to higher-confidence entries with review status of 2-4 stars, ensuring inclusion of SNVs with concordant interpretations from multiple submitters or expert review.

#### Somatic SNVs

Somatic cancer-associated mutations were obtained from the COSMIC ^30,31^. To avoid redundancy arising from recurrent reporting of identical mutations across multiple tumors, COSMIC SNVs were deduplicated at the site level (unique chr:pos:REF:ALT), ensuring a single observation per mutational site.

Only autosomal SNVs were retained in all datasets to ensure comparable genomic coverage across resources.

### Quality control and preprocessing

Quality control procedures were applied to the population-level germline datasets (HGDP and 1KGP). HGDP SNVs were filtered using the PASS flag while excluding SNVs annotated as LOW_VQSLOD or ExcHet. Samples and SNVs with >10% missing genotypes were removed using PLINK2 ^47^. All datasets were represented in the GRCh38 reference assembly, only biallelic SNVs were retained and files were converted to BED format for downstream genomic interval analyses.

### Linkage disequilibrium pruning of germline SNVs

To reduce redundancy arising from haplotype structure, population-level germline SNVs were pruned for LD using PLINK v2 --indep-pairwise 200 25 0.4 command, corresponding to a 200 sliding window, step size of 25 SNVs, and r^2^ threshold of 0.4 ^48^. This procedure retains independent SNVs sites and prevents clustered SNVs within the same haplotype block from inflating overlap counts, for the entire datasets. Because LD reflects inherited structure and does not apply to independently acquired somatic mutations, LD pruning was applied exclusively to HGDP and 1KGP germline datasets (common and rare).

### Allele frequency estimation and SNV stratification

Allele frequencies were calculated separately for each continental superpopulation defined in the datasets using PLINK2 (--freq) ^47^. For HGDP, samples were grouped into the seven geographic regions defined in the original dataset: Africa, America, Central and South Asia, East Asia, Europe, the Middle East, and Oceania ^20^]. For 1KGP, samples were grouped into the five officially defined superpopulations: African (AFR), Admixed American (AMR), East Asian (EAS), European (EUR), and South Asian (SAS) ^33^. Global allele frequencies were also calculated separately for each population reference panel. Common SNVs were defined as variants with global allele frequency ≥1% that also reached allele frequency ≥1% in at least two geographic groups or superpopulations within the corresponding reference panel. SNVs with global allele frequency <1% were classified as rare. Variants with global allele frequency ≥1% that did not meet the two-group criterion were excluded from both frequency classes.

Global allele frequencies were also computed for each reference panel (HGDP or 1KGP). Common polymorphisms (SNPs) were defined as SNVs with global AF ≥1% that were also present at AF ≥1% in at least two superpopulations within the corresponding reference panel. This criterion was adopted to avoid the inclusion of single population-specific mutations, which might be present at high frequency due to evolutionary mechanisms such as genetic drift, founder effects, or inbreeding ^49^. SNVs with global AF <1% were classified as rare.

### Genomic annotation and functional classification

All SNVs were annotated using the Ensembl Variant Effect Predictor (VEP) ^37^. Functional consequences and gene assignments were determined using the MANE (matched annotation from NCBI and EBI) ^38^ transcript set, providing a single representative transcript for each protein-coding gene and minimizing redundancy across isoforms. SNV consequences were summarized according to Ensembl consequence severity categories (including region-based annotations and the most severe coding consequence).

### Exact-coordinate overlap analysis

Overlap between SNV sets was defined as exact genomic coordinate identity (single-base resolution). For *Bedtools* ^50^, each SNV was represented as a 1-bp interval. Because overlap was evaluated at the genomic-position level, each dataset was collapsed to unique genomic sites (chr:pos) prior to analysis, ensuring that recurrent observations or multiple allelic records at the same locus contributed only once to the overlap statistics. Pairwise enrichment of shared loci was assessed using bedtools fisher, which constructs 2×2 contingency tables and performs Fisher’s exact tests using (i) the number of sites in set A, (ii) the number in set B, (iii) the number of overlaps, and (iv) the total genomic search space defined by a genome file. Odds ratios (ORs) were used as effect sizes.

Pairwise enrichment was assessed using bedtools fisher test with default parameters. For WG analyses, the genomic search space corresponded to the combined length of the autosomal chromosomes (GRCh38). For coding-restricted analyses, the background space was defined by the total length of MANE-defined CDS derived from the MANE GFF (General Feature Format) annotation ^38^. WG analyses compared exact genomic sites shared between COSMIC mutations and HGDP common or rare germline SNVs and were independently replicated using the 1KGP. CDS-restricted analyses compared exact genomic sites shared between COSMIC mutations and coding subsets of HGDP common and rare SNVs, as well as ClinVar benign and pathogenic SNV sets.

### Permutation-based null modeling

To independently validate the primary genome-wide enrichment signal, we implemented a chromosome-constrained permutation framework. HGDP common SNV positions were randomly shuffled within their respective autosomes while preserving chromosome assignment, interval length, and the number of SNVs per chromosome. A fixed random seed of 123 was used to ensure reproducibility. Exact-coordinate overlap with COSMIC mutation sites was recalculated for each of 1,000 permutations, generating an empirical null distribution of expected overlap counts. The one-sided empirical P value was calculated using the plus-one correction, P_empirical_ = (*b*+1) / (*B*+1), where *b* is the number of permutations producing an overlap count equal to or greater than the observed value and *B* is the total number of permutations. Because none of the 1,000 permutations produced an overlap equal to or greater than the observed value, the resulting empirical P value was 1/1,001=9.99×10^−4^. Permutation testing was restricted to the primary HGDP common SNV-COSMIC whole-genome comparison; the 1KGP analysis served as an independent population-panel replication, whereas coding-sequence comparisons were evaluated using Fisher’s exact tests within the MANE-defined coding background.

For loci shared between germline and somatic SNV sets, alternate alleles were compared to assess nucleotide-level recurrence. Concordance was summarized as the proportion of shared sites with identical alternate alleles.

### Mutational signature profiling

Shared CDS loci were characterized by mutational signature analysis. Single-base substitution (SBS) signatures were assigned using SigProfilerAssignment ^30,34^, which decomposes observed mutation spectra into known reference signatures. COSMIC v3.4 SBS reference signatures were used to infer the mutational processes contributing to the shared loci ^34–36^.

### Functional enrichment analysis

Genes harboring shared germline-somatic loci were subjected to over-representation analysis (ORA) using the WebGestalt platform ^51^. Each gene was included only once regardless of the number of overlapping SNVs. Enrichment analyses were performed using KEGG (Kyoto Encyclopedia of Genes and Genomes) pathway annotations and cytogenetic band annotations. Statistical significance was assessed using Fisher’s exact test with false discovery rate (FDR < 0.05) correction for multiple testing. ORA output tables were subsequently processed and visualized in R using ggplot2 ^52^.

### Code availability

All code generated for this study is available at: https://github.com/ticatavares/germline_somatic_scripts

