## SUPPLEMENTARY MATERIALS for "Common germline polymorphisms and somatic cancer mutations exhibit non-random positional overlap across the human genome"

**Supplementary Table S1. Summary of input variant records and postprocessed SNV sets used in whole-genome and coding-sequence analyses.** Input values represent the numbers of variant records in the original VCF files before restriction to SNVs. Whole-genome (WG) and coding-sequence (CDS) output values represent the numbers of unique SNV genomic coordinates retained after dataset-specific quality control, selection of biallelic SNVs, allele-frequency classification, and linkage disequilibrium pruning of population-based datasets, where applicable. The primary analyses integrated data from COSMIC, ClinVar, and the Human Genome Diversity Project (HGDP), generating five SNV sets: COSMIC somatic SNVs, ClinVar benign/likely benign SNVs, ClinVar pathogenic/likely pathogenic SNVs, HGDP common SNVs (SNPs), and HGDP rare SNVs (Rares). High-coverage 1000 Genomes Project (1KGP) data provided two additional population-based SNV sets, common and rare, for independent replication of the WG analyses. CDS-restricted analyses were not performed using 1KGP.

| | **HGDP** | | **1KGP** | | **COSMIC** | | **ClinVar** | |
|---|---|---|---|---|---|---|---|---|
| **Input** | 75385302 | | 70692015 | | **Coding** 37261323 | **No coding** 31368583 | 1303558 | |
| **Output** *WG* | **SNPs** 2140477 | **Rares** 13672732 | **SNPs** 1020458 | **Rares** 14191987 | 10587546 | | **Benign** - | **Pathogenic** - |
| **Output** *CDS* | 16040 | 132849 | - | - | 3662887 | | 56794 | 13098 |

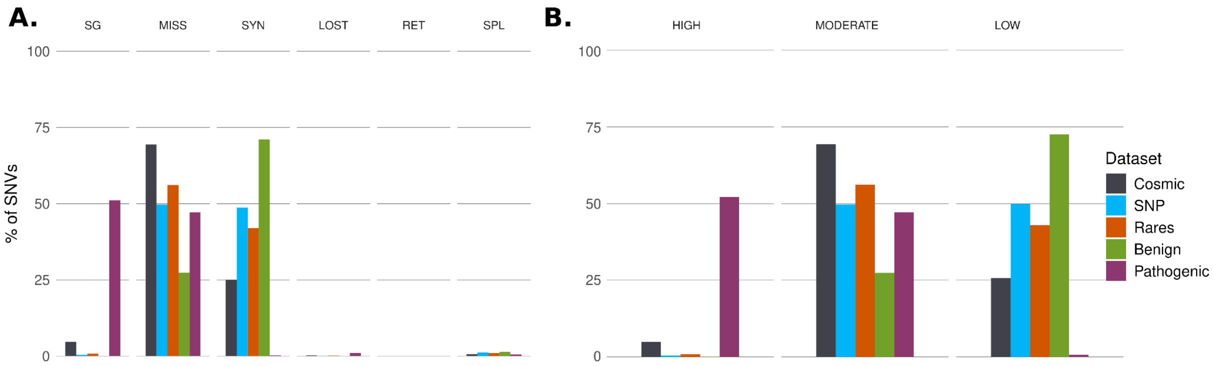


**Supplementary Figure S1: Coding consequences and predicted functional impact of SNVs across the analyzed datasets.** (A) Proportion of SNVs across coding-consequence categories: SG, stop-gained variants, which introduce a premature termination codon and may result in protein truncation or nonsense-mediated decay; MISS, missense variants, which alter the encoded amino acid; SYN, synonymous variants, which do not alter the encoded amino acid; LOST, comprising start-lost variants, which disrupt the translation initiation codon, and stop-lost variants, which abolish the termination codon and may extend the translated protein; RET, stop-retained variants, in which the termination codon is altered but translation termination is preserved; and SPL, splice-region variants, which occur within or near splice sites and may affect pre-mRNA splicing. The figure shows the distribution of these coding consequences across the five primary SNV datasets. (B) Distribution of VEP-predicted impact categories among CDS SNVs across the five datasets: HIGH, variants predicted to have disruptive effects on protein structure or function; MODERATE, variants predicted to alter protein function without necessarily causing complete loss of function; and LOW, variants generally expected to have limited effects on protein function.

**Supplementary Table S2. Complete results of Fisher's exact tests for exact-coordinate overlap analyses.** For each pairwise comparison, the table reports the odds ratio (OR),

95% confidence interval (CI), p-value, and the contingency table generated by bedtools fisher, including the number of overlapping loci (inA_inB), loci unique to each dataset (inA_notB and notA_inB), and the estimated number of loci absent from both datasets (notA_notB). Whole-genome (WG) analyses used the combined length of the autosomal chromosomes (GRCh38) as the genomic search space, whereas coding-sequence (CDS) analyses were restricted to MANE-defined coding regions. The SNP versus Rare comparison was not performed because these datasets are mutually exclusive.

| **WG** | | | | | | | | |
|---|---|---|---|---|---|---|---|---|
| **Comparison** | **OR** | **CI95** | **p_value** | **inA_inB** | **inA_notB** | **notA_inB** | **notA_notB** | **Direction** |
| **HGDP SNPs vs COSMIC** | 1.920 | [1.904, 1.936] | <2.2e-16* | 59569 | 10527977 | 2080908 | 706081926 | Enrichment (OR>1) |
| **HGDP rare vs COSMIC** | 0.668 | [0.664, 0.671] | <2.2e-16* | 136384 | 13536348 | 10451162 | 694626486 | Depletion (OR<1) |
| **1KGP SNPs vs COSMIC** | 2.491 | [2.465-2.517] | <2.2e-16* | 36562 | 10550984 | 983896 | 707178938 | Enrichment (OR>1) |
| **1KGP rare vs COSMIC** | 0.789 | [0.785, 0.793] | <2.2e-16* | 166090 | 10421456 | 14025897 | 694136937 | Depletion (OR<1) |
| **CDS** | | | | | | | | |
| **Comparison** | **OR** | **CI95** | **p_value** | **inA_inB** | **inA_notB** | **notA_inB** | **notA_notB** | **Direction** |
| **SNPs vs COSMIC** | 5.652 | [5.476, 5.833] | <2.2e-16* | 9575 | 6465 | 3653035 | 13940452 | Enrichment (OR>1) |
| **Rare vs COSMIC** | 0.522 | [0.516, 0.528] | <2.2e-16* | 39337 | 93512 | 3623529 | 4493760 | Depletion (OR<1) |
| **COSMIC vs Benign** | 0.699 | [0.687, 0.711] | <2.2e-16* | 20372 | 3642515 | 36422 | 4550829 | Depletion (OR<1) |
| **COSMIC vs Pathogenic** | 0.557 | [0.537, 0.578] | 1.1E-225* | 4036 | 3658851 | 9062 | 4578189 | Depletion (OR<1) |
| **SNPs vs Benign** | 66.480 | [63.743, 69.335] | <2.2e-16* | 2728 | 13312 | 54066 | 17539421 | Enrichment (OR>1) |
| **SNPs vs Pathogenic** | 0.335 | [0.126, 0.893] | 1.3E-02 | 4 | 16036 | 13094 | 17580393 | Depletion (OR<1) |
| **Rare vs Benign** | 11.153 | [10.889, 11.423] | <2.2e-16* | 8339 | 124510 | 48455 | 8068834 | Enrichment (OR>1) |
| **Rare vs Pathogenic** | 0.267 | [0.206, 0.346] | 2.6E-02 | 57 | 132792 | 13041 | 8104248 | Depletion (OR<1) |
| **Benign vs Pathogenic** | 2.150 | [1.864, 2.480] | <2.2e-16* | 192 | 56602 | 12906 | 8180438 | Enrichment (OR>1) |

**bedtools reports P = 0 because of floating-point precision; values are shown as $P < 2.2 \times 10^{-16}$*

**Supplementary Table S3. Enrichment of canonical mutational hotspots among shared germline-somatic loci.** Fisher's exact tests assessed the overlap of the 59,569 exact-coordinate loci shared between HGDP common germline SNVs and COSMIC somatic mutations with CpG islands and microsatellite regions. CpG islands intersected 2,839 shared loci (4.77%; OR = 7.49), whereas microsatellites intersected 131 shared loci (0.22%; OR = 3.85). Although both contexts were significantly enriched, they encompassed only small proportions of the complete set of shared loci.

| Dataset A | Dataset B | *p* | OR | in_A/in_B | in_A/not_B | not_A/in_B | not_A/not_B |
|---|---|---|---|---|---|---|---|
| Shared germline-somatic loci | CpG Island | 0 | 7.49 | 2839 | 56730 | 24027 | 3595100 |
| Shared germline-somatic loci | Microsatellite | 4e-37 | 3.85 | 131 | 59438 | 37753 | 65914793 |

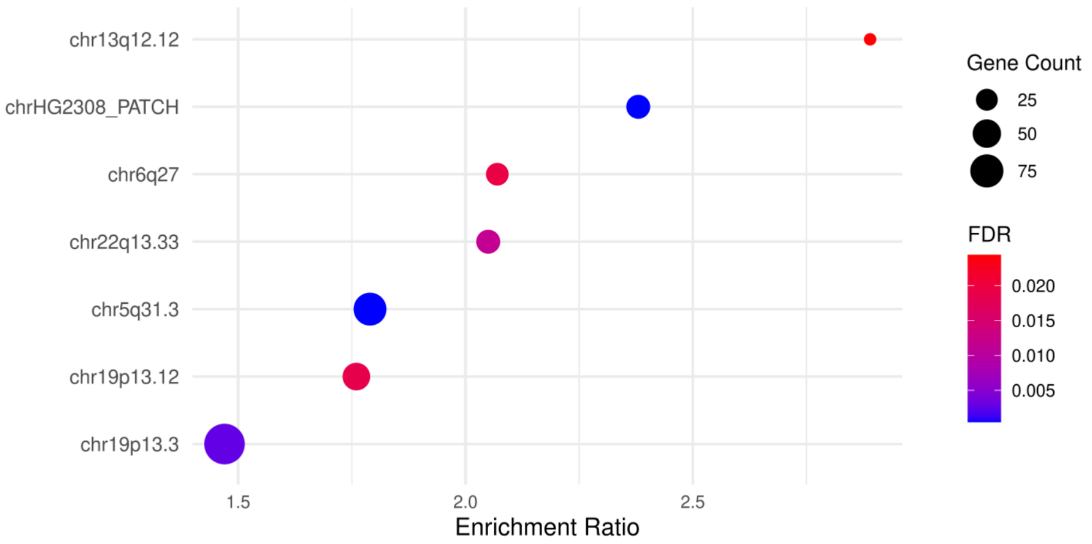


**Supplementary Figure S2. Cytogenetic band enrichment analysis.** The Y-axis represents significantly enriched genomic regions (FDR < 0.05), whereas the X-axis shows the enrichment ratio. The dot size reflects the number of genes within each region, and the color scale represents statistical significance, ranging from blue (most significant) to red (less significant).

**Supplementary Table S4. Complete list of genes enriched in the KEGG pathways identified in the Over-Representation Analysis (ORA).** The table presents all significantly enriched pathways (FDR < 0.05), along with the mapped genes associated with each pathway.
EXCEL FILE

**Supplementary Table S5. Complete list of genes present in the significantly enriched cytogenetic bands identified in the Over-Representation Analysis (ORA).** The table includes all cytogenetic bands with FDR < 0.05, along with their respective mapped genes.
EXCEL FILE
